# Evaluating Climate Warming Effects on Soil Resistome and Pathogenome: Future Risks for Agriculture and Human Health

**DOI:** 10.1101/2024.01.22.576767

**Authors:** Zhiguo Zhang, Feng Ju

## Abstract

How climate change affects soil antibiotic resistome (i.e., the collection of antibiotic resistance genes) is a critical question for environmental and human health. By examining the dynamics of soil resisomes in a six-year (2014-2019) climate change experiment, this study provides explicit insights into the risk of antibiotic resistance in cropland and grassland microbiomes under future climate scenarios. Extreme summers (+2.2°C during 2018-2019) significantly shifted the resistomic composition, resulting in a prominent increase in abundance of ARGs (copy/cell) conferring resistance to novobiocin (52.7%-72.8%), tetracycline (32.5%-53.0%) and vancomycin (31.5%-62.9%). Importantly, simulated warming (+0.6°C) significantly increased the proportion of mobilizable ARGs, possibly resulting from the SOS response of soil microbes stimulated by warming-induced drought. In contrast, extreme summers decreased the mobility potential by dramatically filtering the hosts (e.g., *γ-Proteobacteria*) of mobilizable ARGs. Climate warming and extreme summers also offer a worrisome competitive advantage for specific soil-dwelling antibiotic-resistant phytopathogens (*Clavibacter michiganensis* and *Rhodococcus fascians*) and human pathogens (e.g., *Staphylococcus aureus* and *Mycobacterium tuberculosis*), which escalates the risk of outbreaks for specific plant and human infectious diseases. Overall, our findings emphasize the urgent need for continuous monitoring of soil ARGs and pathogens under the on-going global change. Such efforts are crucial for safeguarding human health and ensuring the sustainability of modern agriculture within a global One-Health framework.

**Synopsis:** Climate warming increases the mobility potential of soil antibiotic resistance genes, and both climate warming and extreme summers escalate the risk of outbreaks for specific plant and human infectious diseases.

## Introduction

Antibiotic resistance has become a global health challenge owing to its continuous emergence and rapid dissemination between microbes in humans, animals, and the environment ^1^. It is estimated that antibiotic resistance could claim 10 million annual deaths and result in a cumulative economic loss of 100 trillion US dollars by 2050 ^2^. This impending crisis is being aggravated in the face of global climate warming ^3^. Antimicrobial surveillance suggested that elevated ambient temperature was significantly associated positively with the antibiotic resistance prevalence in clinical isolates of human bacterial pathogens, such as *Klebsiella pneumoniae* and *Pseudomonas aeruginosa* ^4-6^, which may be attributed to the accelerated rate of horizontal gene transfer ^4^. Higher antibiotic resistance levels of aquaculture-related bacterial pathogens were also observed to be correlated with warmer temperatures ^7^. These results suggest that climate warming potentially promotes the development of antibiotic resistance in bacterial pathogens.

Despite the increasingly noticeable impacts of elevated ambient temperature on antibiotic resistance prevalence in numerated bacterial isolates, whether and how climate warming affects soil resistome and antibiotic-resistant pathogens in natural environments has not yet been clearly known. Based on limited reports, simulated warming remarkably increased antibiotic resistance gene (ARG) abundance in river water ^8^. Also, climate warming tended to increase the proportion of specific ARGs in natural forest soils ^9^. However, the impacts of long-term experimental warming on soil resistomes, regarding ARG mobility and host pathogenicity, which are key traits associated with the evolution, dissemination and health risk of ARGs ^10,11^, are poorly understood.

To address the above-mentioned research gaps, we examined the ARG diversity, composition, mobility potential and host pathogenicity of cropland and grassland soil microbiomes over 6 years from 2014 to 2019. We achieved this goal by mining a publicly available data set of shotgun metagenomes generated from the experiments at the Global Change Experimental Facility (GCEF) research station in Central Germany ^12,13^, wherein climate conditions were manipulated to simulate projected climatic scenarios spanning the period from 2070 to 2100 ^13^. In addition, Central Europe underwent an extraordinary drought starting in the spring of 2018 and persisting until the fall of 2019. This unforeseen event provided a distinctive opportunity to explore the effects of extreme summers on soil resistomes. This study aims to address (i) whether and how experimental warming and extreme summers affect soil ARG diversity, composition, and mobility; (ii) whether and how experimental warming and extreme summers affect soil-borne antibiotic-resistant pathogens; and (iii) what are their underlying mechanisms. Overall, this study delivers crucial insights into the response of soil resistome, such as ARG mobility and host pathogenicity, to future climate scenarios in cropland and grassland soils.

## Materials and methods

### Data collection from samples of the Global Change Experimental Facility

The publicly available metagenomic datasets of soil microbiomes from Bei et al. ^13^ were utilized to explore the impacts of future climate scenarios on soil resistome. In brief, 180 samples (2 climates × 3 land-use types × 5 replicates × 6 summers) were collected from conventional cropland (CC), organic cropland (OC), and intensive grassland (IG) in the Global Change Experimental Facility (GCEF) during 2014-2019. This facility consists of 10 blocks with 5 plots each, resulting in a total of 50 plots (each 16m × 24 m). Half of the plots were exposed to simulated warming (+ 0.6 °C), while the other half were subjected to ambient climate conditions. Detailed information on the experiment could be seen in the previous study ^13^. In total, 8.64 Tb of FASTQ reads were generated from 180 metagenomes, with an average of 48 Gb per soil sample. The metagenomic sequencing data are accessible in NCBI with the accession number PRJNA 838942.

### Metagenomic read quantification of ARGs, MGEs and pathogen-host interaction (PHI) genes

For each metagenome analyzed in this study, raw sequencing reads were processed with Fastp (v0.23.1) ^14^ to remove Illumina adaptors, low-quality reads, and duplicate reads. The generated clean reads of each metagenome were used to annotate and quantify ARGs using the ARGs-OAP pipeline (v3.2.2) ^15^. To eliminate the bias resulting from different sequencing depths, each metagenome was randomly subsampled to an even depth of 106 million clean reads using Seqtk (v1.4, https://github.com/lh3/seqtk). The unit of gene copy per cell (GPC) was used to present the relative abundance of ARGs ^10^. This unit normalizes sequencing depth, ARG lengths, and prokaryotic cells or genomes in the metagenome and has a straightforward biological and clinical meaning ^16^. The ARG diversity was calculated as the number of ARG subtypes detected in each metagenome. The ARG-assigned reads were further taxonomically annotated in order to identify microbial hosts of ARGs using Kraken2 (v2.1.3) with default settings ^17^. The identification of pathogenic species was based on the taxonomic matching with the pathogen list curated by the previous study ^18^.

The process of MGE quantification was performed using a customized Local Analysis Pipeline of MGEs (MGEs-LAP, Fig. S1) that inherits the two-step search strategy of ARGs-OAP pipeline ^15^ to eliminate false-positive hits in mobileOG-db ^19^, a manually curated database of MGEs. Briefly, the clean reads of each metagenome were first filtered by aligning them to the database using Diamond with a loose cutoff (identity > 60%, coverage > 15%, *E* value < 10). Then, the filtered reads were searched against the database again to identify MGE-like reads using BLAST with a strict cutoff (identity > 80%, coverage > 75%, *E* value < 1e-7). The abundance (GPC) of each MGE gene in one specific metagenome was calculated with the following equation:

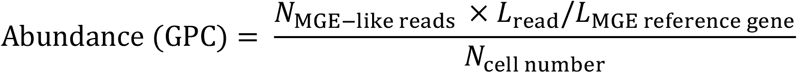

Where *N*_MGE-like reads_ is the number of MGE-like reads annotated to one specific MGE gene; *L*_read_ is the read length; *L*_MGE reference gene_ is the nucleotide length of the corresponding MGE gene; and *N*_cell number_ is the estimated cell number of one specific metagenome.

The method of PHI quantification is similar to MGE quantification, except that the database was changed to the pathogen-host interactions (PHI) database (v4.16) ^20^. This database compiles expertly curated molecular and biological information on experimentally verified pathogenicity, virulence, and effector genes demonstrated to affect and indicate the outcome of pathogen-host interactions ^20^.

### Metagenome assembly and binning and metagenome-assembled genome (MAG) analysis

The clean reads of the aforementioned soil metagenomes for the replicated samples were combined, which resulted in 36 metagenomes representing 2 climates, 3 land-use types and 6 summers. For each combined metagenome, clean reads were *de novo* assembled using Megahit (v1.2.9) ^21^. The metagenome assembly results were summarized in Dataset S1. The coverage profiles of each assembly were generated by mapping clean reads from 12 combined metagenomes with the same land use using BWA ^22^. The scaffolds of each assembly were subsequently clustered into genome bins informed by the coverage profiles using MetaBAT2 ^23^ with default parameters. This recently proposed cross-sample binning strategy is demonstrated to significantly improve the binning yields of soil metagenomes, as validated by our prior study ^24^. The bins with an overall quality of > 50% (completeness – 5 × contamination) were considered eligible ones ^25^. Finally, 820 eligible bins were obtained (Dataset S2). Furthermore, the bins were dereplicated using dRep (v3.0.0) ^26^ with a 95% ANI threshold, finally resulting in 130 species-level representative MAGs. MAGs were taxonomically assigned using the classify_wf module of gtdbtk (v2.1.1) ^27^ and were classified into 122 bacterial MAGs and 8 archaeal MAGs. Phylogenetic analysis of bacterial MAGs was conducted with the gtdbtk infer module based on a set of 120 bacteria-specific marker genes from GTDB ^27^, and the phylogenetic trees were visualized in iTOL ^28^. The abundance of MAGs in each sample was quantified using CoverM (v0.6.1) (https://github.com/wwood/CoverM).

### Functional annotation and quantification of metagenome-assembled ARGs, MGEs, and plasmids

Open reading frames (ORFs) were predicted for each contig using Prodigal (v2.6.3) ^29^. The ARG-like ORFs were identified by aligning their nucleotide sequences to the SARG database (v3.2.2) ^15^ using BLASTX at an E value ≤ 10^−7^ with at least 60% identity and 80% query coverage. The ORFs located on ARG-carrying contigs were then annotated against the mobileOG-db database ^19^ using BLASTX with the above threshold. The ARGs were considered to have mobility potentials if their distance with MGEs is less than 10 kb ^30, 31^. Furthermore, the mobility potential of an ARG cassette was quantified by mobility incidence (M%) as proposed by Ju et al., 2019 ^32^. The plasmid sequences of ARG-carrying contigs were predicted using PlasClass (v0.1.1) ^33^ with default parameters. The taxonomy information of ARG-carrying contigs was annotated using CAT (v5.3) ^34^ with default settings. The abundance of ARG-carrying contigs in each sample was quantified using CoverM (v0.6.1).

### Statistical analysis

Linear mixed-effects models (LMMs) were used to evaluate the effects of simulated warming and extreme summers on the ARG diversity and the abundance of pathogens. The lme4 (v1.1.34) ^35^ package was used to implement LMMs in R. In the test of simulated warming, simulated warming (0 for ambient temperature and 1 for simulated warming) treatments were considered as fixed effects while sampling time (year) and block were termed as random intercept effects (y ∼ simulated warming + (1 | year) + (1 | block)). In the test of extreme summers, extreme summers (0 for normal summers and 1 for extreme summers) were considered as fixed effects, while block was termed as random intercept effects (y ∼ extreme summers + (1 | block)). Effect sizes of treatments were represented by the regression coefficients in the LMMs. Wald type II χ^2^ tests were used to calculate the *p* values from the LMMs using the car package ^36^.

DESeq2 (v1.40.2) ^37^ was used to identify significant changes in the abundance of ARGs and MAGs in response to simulated warming and extreme summers with the formula (design= ∼year + land use + simulated warming or design= ∼land use + extreme summers). The *p* values were adjusted for multiple testing using the Benjamini and Hochberg (BH) method.

## Results

### Simulated warming and extreme summers reduce soil ARG diversity and reshape soil resistome

Metagenomic read-based analysis detected a total of 929 ARG subtypes encoding resistance to 26 different classes of antibiotics in 180 soil metagenomes from three land use types, i.e., conventional cropland (CC), organic cropland (OC) and intensive grassland (IG). The total abundance and richness of ARGs exhibited ranges of 0.104 to 0.187 GPC and 166 to 304 subtypes in CC soils, 0.098 to 0.175 GPC and 172 to 227 subtypes in OC soils, and 0.118 to 0.212 GPC and 176 to 295 subtypes in IC soils. Statistical analysis showed that the ARG abundance and richness in CC (with herbicides and pesticides applied) were significantly higher than those of OC by an average of 5.0% and 9.3%, respectively (Wilcoxon Signed-Rank test, *p* < 0.01; Fig. S2). This result indicates that herbicides and pesticides may promote the dissemination of soil ARGs.

Further analysis showed that simulated warming (+0.6°C and -9.1% soil moisture) significantly decreased the ARG richness of CC by 4.4% (LMMs, *p* < 0.01) and a marginally significant decrease in ARG richness of IG soils by 4.2% (LMMs, *p* < 0.07). Moreover, extreme summers (+2.2°C and -35.4% soil moisture during 2018-2019) significantly reduced ARG richness in CC, OC and IG soils by 12.5%, 6.7% and 7.8%, respectively (LMMs, *p* < 0.001; Fig. 1a). Specifically, the reduced richness was mainly attributed to aminoglycoside, beta-lactam, multidrug, polymyxin and trimethoprim resistance genes (Fig. S3). In contrast, the total ARG abundance almost showed no significant response to simulated warming and extreme summers (Fig. 1b) except for the significant influence noticed in CC and IG soils by extreme summers (DESeq2, *p* < 0.05) and simulated warming (DESeq2, *p* < 0.001), respectively (Fig. 1b).

**Figure 1.**
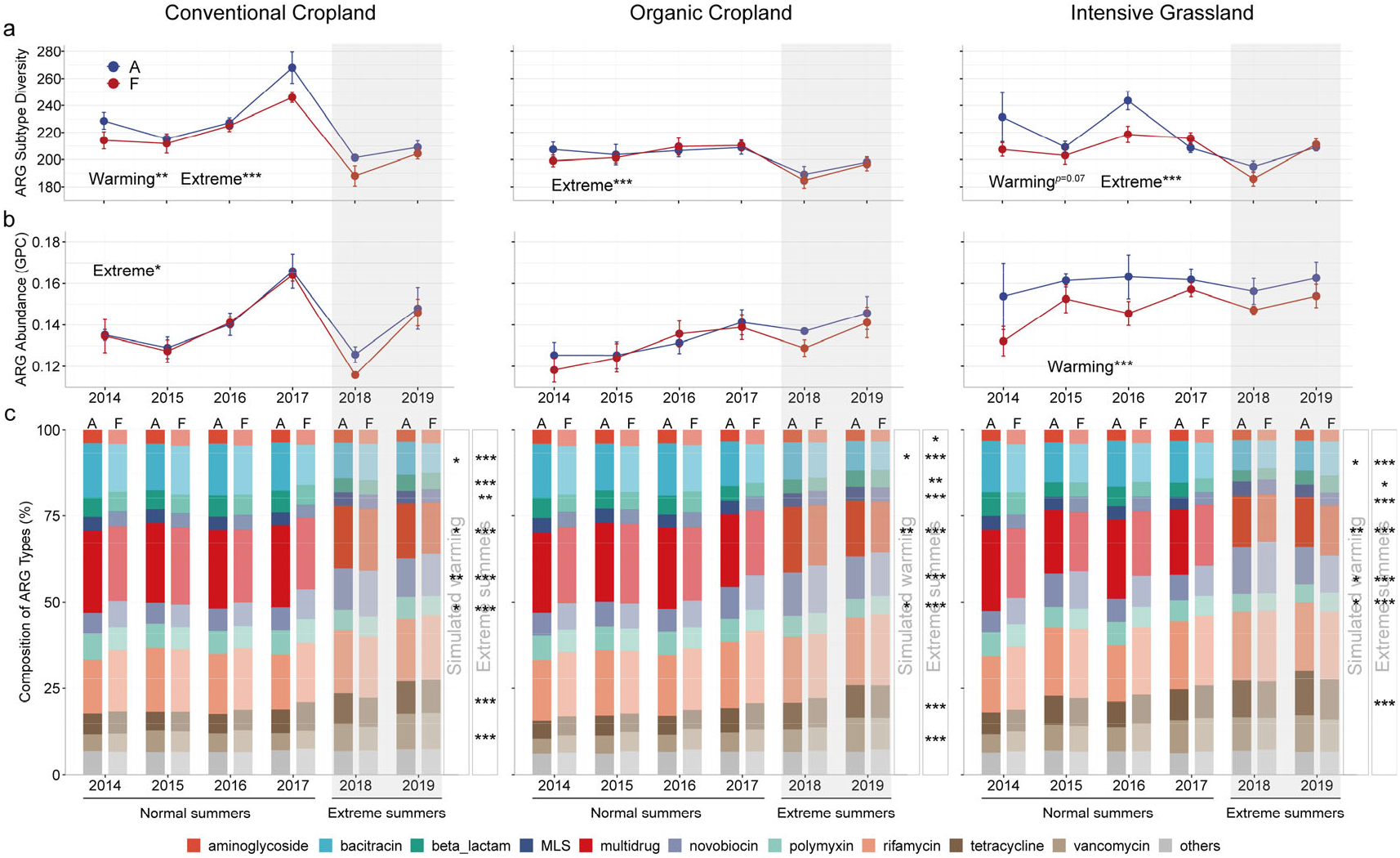
The variation of resistome features under simulated warming and extreme summers. **a**. The ARG richness in three land-use types. Statistical significance was determined using linear mixed-effects models and Wald type II χ^2^ tests (n = 60). **b**. The ARG abundance in three land-use types. Statistical significance was determined using DESeq2. **c**. The proportions of ARG types in three land-use types. Statistical significance was determined using DESeq2. \*\*\**p* < 0.001, \*\**p* < 0.01, \**p* < 0.05. A: ambient; F: future.

Extreme summers significantly impacted the composition of soil ARGs, resulting in a decreased abundance of bacitracin (by 25.9%-34.9%), beta-lactam (10.7%-30.4%), multidrug (18.4%-30.3%) and polymyxin (9.1%-20.7%) resistance genes and an increased abundance of novobiocin (52.7%-72.8%), tetracycline (32.5%-53.0%) and vancomycin (31.5%-62.9%) resistance genes (DESeq2, *p* < 0.001-0.05; Fig. 1c). PCoA analysis also revealed a significant separation of ARG composition between normal and extreme summers (ANOSIM, *R*^2^ = 0.23, *P* = 0.001; Fig. S4). In contrast, simulated warming caused a lesser shift in ARG composition (ANOSIM, *R*^2^ = 0.02, *P* < 0.01), of which bacitracin (8.3%-13.5%), multidrug (8.8%-16.9%) and polymyxin (6.5%-12.0%) resistance genes significantly decreased while novobiocin (4.8%-10.9%) resistance genes increased (DESeq2, *p* < 0.01-0.05; Fig. 1c).

### Simulated warming and extreme summers exhibit opposite impacts on soil ARG mobility

To investigate the mobility potential of ARGs under simulated warming and extreme summers, ARGs and mobile gene elements (MGEs) located on the metagenome-assembled contigs were annotated. In total, we detected 51,508 ARG sequences located on 50,051 contigs, of which 478 ARGs were colocalized (distance < 10 kb) with diverse MGEs (Dataset S3). Furthermore, 6320 (12.6%) ARG-carrying contigs were identified as plasmid sequences. Of note, most of these ARG-carrying plasmids were taxonomically assigned to *Proteobacteria* (35.5%) and *Actinomycetota* (25.1%) (Dataset S3).

We found that extreme summers and simulated warming influenced the ARG mobility of soil microbiome in intriguingly opposite ways. Simulated warming significantly increased the proportion of ARG-carrying contigs flanked with MGEs by 1.7% (Wilcoxon Signed-Rank test, *p* < 0.05; Fig. 2a), indicating that climate warming will increase the mobility potential of ARGs. Whereas, extreme summers significantly decreased the proportion of ARG-carrying contigs flanked with MGEs by 9.8% (Mann-Whitney test, *p* < 0.001; Fig. 2a). For specific MGE types and land-use types, simulated warming remarkably increased the proportion of ARG-carrying contigs flanked with integration elements by 17.2%, 17.9% and 6.1% in CC (*p* < 0.05), OC (*p* < 0.05) and IG (*p* = 0.14) soils, respectively (Wilcoxon Signed-Rank test, Fig. 2c-e), while extreme summers significantly decreased the proportion of ARG-carrying contigs with phage elements (by 7.6%-14.9%, *p* < 0.001-0.05) and recombination elements (by 18.4%-31.9%, *p* < 0.001-0.01) in three land-use types (Mann-Whitney test, Fig. 2c-e).

**Figure 2.**
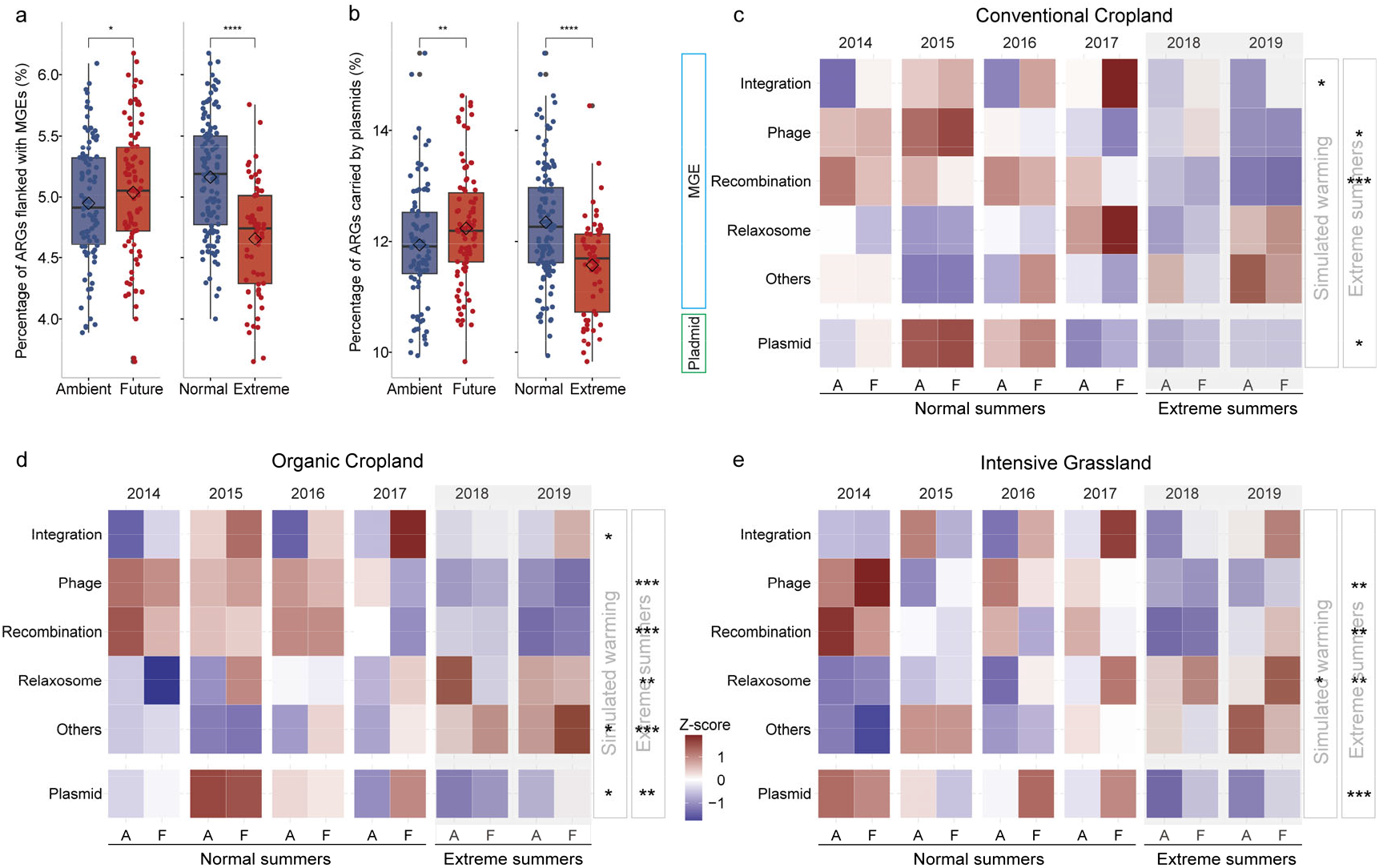
The mobility potential of ARGs under simulated warming and extreme summers. **a**. The proportion of ARGs flanked with MGEs under simulated warming and extreme summers. **b**. The proportion of plasmid-borne ARGs under simulated warming and extreme summers. **c, d, e**. The proportion of ARGs flanked with different MGE types under simulated warming and extreme summers in conventional cropland, organic cropland and intensive grassland. Significant effects of simulated warming and extreme summers are based on Wilcoxon Signed-Rank test and Mann-Whitney test, respectively. *p* values were adjusted for multiple testing using the Benjamini-Hochberg (BH) method; \*\*\**p* < 0.001, \*\**p* < 0.01, \**p* < 0.05. The proportion of genes was row-scaled.

Echoing the impacts on ARG-MGE cooccurrence patterns, our results also showed that the proportion of ARG-carrying plasmids significantly increased by 2.5% under simulated warming (Wilcoxon Signed-Rank test, *p* < 0.01) but decreased by 6.3% under extreme summers (Mann-Whitney test, *p* < 0.001; Fig. 2b). For specific land-use types, extreme summers significantly reduced the proportion of ARG-carrying plasmids by 5.8%, 6.1% and 7.1% in CC, OC and IG soils, respectively (Mann-Whitney test, *p* < 0.001-0.05; Fig. 2c-e), revealing the consistent impacts of extreme summers on soil resistome cross different land-use types.

### Future climate scenarios shift the host identity of soil ARGs, especially for multi-antibiotic-resistant pathogens

To explore the changes of ARG hosts under future climate scenarios (i.e., simulated warming and extreme summers), a total of 122 species-level bacterial MAGs were obtained, which were affiliated to 12 phyla, such as *Proteobacteria* (30) and *Actinomycetota* (27), *Acidobacteriota* (15) and *Bacteroidota* (15) (Fig. 3a). Among these bacterial MAGs, 96 were found to carry at least one ARG. Notably, *γ-Proteobacteria* harbored the largest number of ARG-carrying MAGs (each containing 6.95 ARGs on average), followed by *Actinomycetota* (Fig. 3a).

**Figure 3.**
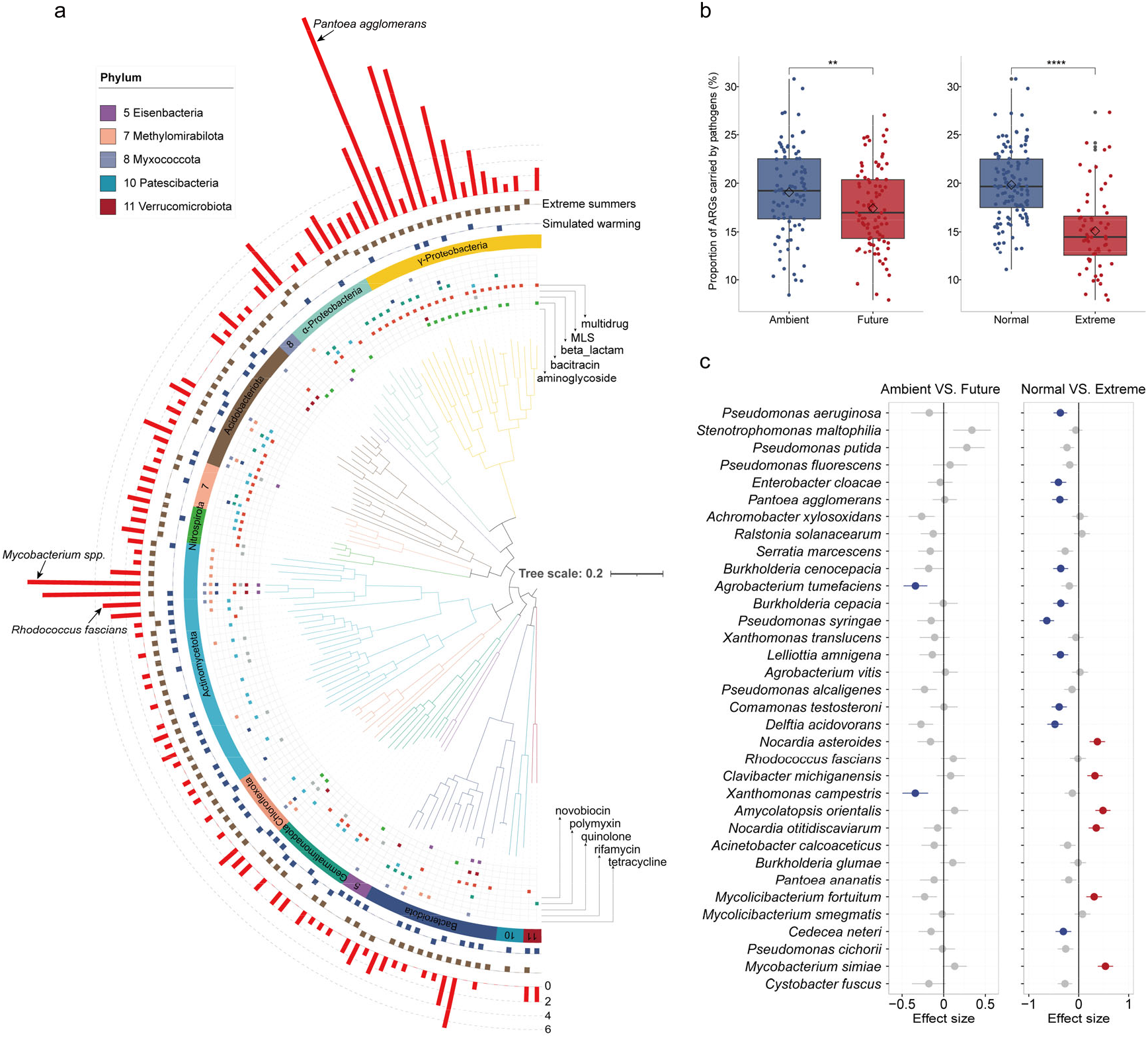
Phylogenomic distribution of ARG-carrying bacteria in response to simulated warming and extreme summers. **a**. Phylogenetic distribution of 122 species-level representative bacterial MAGs. The phylogenetic tree was constructed based on 120 bacterial marker genes using GTDBT. The red bar shows the maximum amounts of ARGs in each genome cluster (95% ANI). The brown and blue bars indicate whether the genomes were enriched under extreme summers and simulated warming, respectively. The square symbol with different colors indicates the ARG types in each genome. **b**. The proportion of ARGs carried by human/animal/plant pathogens under simulated warming and extreme summers. **c**. Effect sizes of simulated warming (left) and extreme summers (right) on the (rescaled) abundance of ARGs carried by human/animal/plant pathogens (detected in ≥ 10 samples) based on linear mixed-effects models. Data are presented as mean ± s.e.m. of the estimated effect sizes. Statistical significance is based on Wald type II χ^2^ tests (n = 180). Significantly positive and negative changes are denoted by red and blue dots, respectively; Non-significant change is denoted by grey dots.

Of the 96 ARG-carrying MAGs, 54 showed significant abundance changes under simulated warming (Fig. 3a). Specifically, 29 MAGs (e.g., *Actinomycetoa* and *Chloroflexota*) experienced an increased abundance, whereas 25 MAGs (e.g., *γ-Proteobacteria, Bacteroidota* and *Acidobacteriota*) showed decreased abundance (DESeq2, *p* < 0.0001-0.05; Fig. 3a). During extreme summers, 79 MAGs displayed significant alterations, with 29 MAGs (e.g., *Actinomycetoa, α-Proteobacteria* and *Chloroflexota*) experiencing an increased abundance and 50 MAGs (e.g., *γ-Proteobacteria, Bacteroidota, Gemmatimonadota* and *Acidobacteriota*) showing decreased abundance (DESeq2, *p* < 0.0001-0.05; Fig. 3a). In general, both simulated warming and extreme summers significantly reduced abundance of ARG-carrying *γ-Proteobacteria*, while both conditions concurrently increased abundance of ARG-carrying *Actinomycetota*. For example, extreme summers witnessed a 63.9% decrease in abundance of *Pantoea agglomerans*, an uncommon but multi-antibiotic-resistant *γ-Proteobacteria* (29 ARGs), whereas a 39.7% increase in abundance of highly antibiotic-resistant *Mycobacterium spp.* of *Actinomycetota* (15 ARGs) (Fig. 3a). In particular, one phytopathogen *Rhodococcus fascians* (5 ARGs) significantly increased by 20.9% and 8.2% during simulated warming and extreme summers, respectively (Fig. 3a).

To further disentangle the dynamics of ARG-carrying pathogens under future climate scenarios, ARG-like reads were taxonomically annotated to identify their microbial hosts. Subsequently, 34 human/animal/plant pathogens (detected in ≥ 10 samples) were discerned based on the derived taxonomic information (Dataset S4). The results showed that 15.1% of ARG-like reads were annotated as human/animal/plant pathogens (Fig. 3b). In contrast, only 6.7% of total metagenomic reads were annotated as human/animal/plant pathogens (Fig. S5), indicating the marked enrichment of ARGs in pathogens. Moreover, simulated warming and extreme summers significantly decreased the proportion of pathogen-carrying ARGs by 8.9% (Wilcoxon Signed-Rank test, *p* < 0.01) and 23.9% (Mann-Whitney test, *p* < 0.0001), respectively (Fig. 3b). Specifically, only two pathogenic species (i.e., *Agrobacterium tumefaciens* and *Xanthomonas campestris*) showed significant response to simulated warming (LMMs, *p* < 0.05; Fig. 3c, left panel). In contrast, 16 of the 34 pathogenic species showed significant abundance changes under extreme summers (LMMs, *p* < 0.0001-0.05; Fig. 3c, right panel). Interestingly, all 10 of the decreased species under extreme summers belong to β- or γ-*Proteobacteria*. In contrast, the 6 increased species belong to *Actinomycetota*, such as the phytopathogen *Clavibacter michiganensis* and the human pathogen *Nocardia asteroides*. These results together indicate that future climate scenarios will reshape the host composition of soil ARGs and facilitate the prevalence of specific antibiotic-resistant pathogens.

### Future climate scenarios reshape soil pathogenome and selectively enrich specific soil-borne pathogens

To further validate the above-discovered shift in soil pathogen composition under future climate scenarios, we also annotated and quantified soil-dwelling pathogens using the pathogen-host interactions (PHI) database (v4.16) ^20^. This database compiles expertly curated molecular and biological information on genes demonstrated to affect the outcome of pathogen-host interactions. Of the 285 pathogenic species in the database, 244 species were detected in soil metagenomes of this study, including bacterial, fungal and protistan pathogens that cause various diseases in humans, animals, and crops (Dataset S5).

Consistent with the aforementioned overall decrease in ARG-carrying pathogens (Fig. 3b), simulated warming and extreme summers significantly decreased the total PHI abundance by 4.1% and 12.5%, respectively (*p* < 0.0001, Fig. 4a). For the most abundant 30 species (0.03-0.42 GPC), most of the species significantly decreased under simulated warming (LMMs, *p* < 0.0001-0.05; Fig. 4b) or extreme summers (LMMs, *p* < 0.0001-0.05; Fig. 4c). These decreased species mainly belong to *Proteobacteria* and *Ascomycota*. Notably, four human bacterial pathogens, i.e., *Staphylococcus aureus, Mycobacterium tuberculosis, Corynebacterium diphtheriae* and *Borreliella burgdorferi*, showed an increasing trend under simulated warming (Fig. 4b) and significantly increased by 2.1%-26.1% under extreme summers (LMMs, *p* < 0.0001-0.01; Fig. 4c). In addition, we also detected 16 species with lower abundance (< 0.03 GPC) that significantly enriched under extreme summers (LMMs, *p* < 0.0001-0.05). These species included crop pathogens such as *Verticillium dahliae* and *Clavibacter michiganensis*, human pathogens like *Streptococcus pneumoniae* and *Listeria monocytogenes*, as well as the pig pathogen *Brucella suis* (Fig. S6).

**Figure 4.**
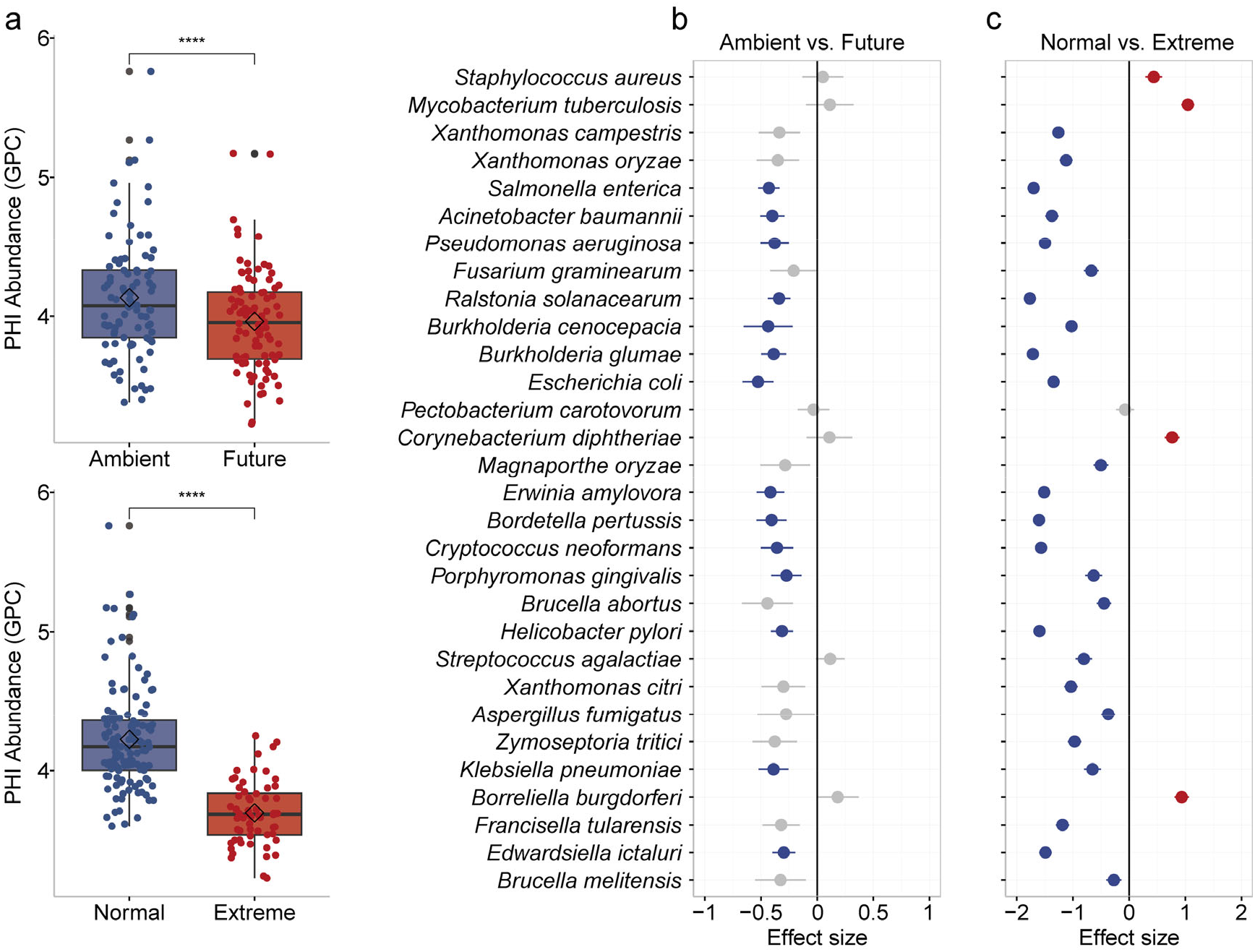
The variation of pathogen-host interactions (PHI) related genes and their hosts in response to simulated warming and extreme summers. **a**. The total abundance of PHI genes under simulated warming and extreme summers. Significant effects of simulated warming and extreme summers are based on Wilcoxon Signed-Rank test and Mann-Whitney test, respectively. **b, c**. Effect sizes of simulated warming and extreme summers on the (rescaled) abundance of top-30 PHI hosts based on linear mixed-effects models. Data are presented as mean ± s.e.m. of the estimated effect sizes. Statistical significance is based on Wald type II χ^2^ tests (n = 180). Significantly positive and negative changes are denoted by red and blue dots, respectively; Non-significant change is denoted by grey dots.

## Discussion

Understanding how climate change affects soil resistomes and the underlying mechanisms is a critical issue for the environment and human health. By examining the dynamic changes of soil resistome in a consecutive six-year (2014-2019) climate change experiment, this study provides important insights into the risk of antibiotic resistance in cropland and grassland soils under future climate scenarios related to the loss of ARG diversity, alteration in resistomic composition, change of ARG mobility, and shift in soil-borne pathogenome.

Both climate warming and extreme summers are found to consistently reduce the ARG diversity and alter the ARG composition across different years. The decreased ARG diversity may result from the decrease in microbial diversity (Fig. S7). The warming conditions will induce more intense competition among soil microbes ^38^, eventually leading to various microbial extinction events and ultimate biodiversity decrease ^39^. During extreme summers (2018-2019), the soil moisture decreased by 35.4%-56.4%. The reduced moisture would act as a strong filtering factor against existing microbial species, which could also cause soil biodiversity loss ^40, 41^. As a result, some subtypes of ARGs, such as aminoglycoside-resistant *APH(3’)-IIc* and carbapenem-resistant *IMP-18* in this study, are expected to disappear along with their microbial hosts.

The alteration of microbial composition also leads to changes in ARG composition and mobility potential. Our results revealed a significant increase in the abundance of *Actinomycetota*, the dominant hosts of Novobiocin, tetracycline and vancomycin resistance genes, coupled with a significant reduction in the abundance of *Proteobacteria*, the dominant hosts of bacitracin, beta-lactam, multidrug, and polymyxin resistance genes, under simulated warming. These changes were much more pronounced during extreme summers. Therefore, the ARG composition was significantly changed under simulated warming and extreme summers. It is reasonable to project that future climate scenarios will increase the prevalence and health risk of specific types of ARGs, such as tetracycline and vancomycin resistance genes, which may transfer from soil-dwelling microbes to clinical-prevalent pathogens. For example, exploring a 30,000-year-old Alaskan permafrost soil found that a complete *vanHAX* vancomycin resistance operon is similar to current clinical ARGs ^42^.

Interestingly, simulated warming significantly increased the mobility potential of ARGs, whereas extreme summers decreased the mobility potential of ARGs. It was proposed that elevated temperature may facilitate the horizontal transfer of ARGs, which could lead to increased antibiotic resistance in clinical pathogenic isolates ^4^. In addition, drought impacts a suite of physiological processes in microbes by elevating levels of damaging reactive oxygen species and inducing DNA strand breaks, therefore triggering the up-regulation of the SOS response ^43^. The SOS response could, in turn, remarkably promote the horizontal transfer of ARGs ^44, 45^. Therefore, warming-induced precipitation changes (-9.1% soil moisture) may trigger the SOS response of soil bacteria, leading to the increased HGT frequency of mobilizable ARGs under simulated warming. Compared with simulated warming, extreme summers significantly eliminated *Proteobacteria* (by 14.9%), the dominant hosts of mobilizable ARGs (Dataset S3). Therefore, extreme summers greatly decreased the abundance of hosts of mobilizable ARG, leading to the decrease of ARG mobility potential.

Strikingly, specific ARG-carrying pathogens and soil-borne pathogens significantly increased under simulated warming and extreme summer, which mainly phylogenetically belong to gram-positive species. A thick peptidoglycan cell wall and sporulation can assist bacteria, especially gram-positive species, in tolerating drought ^46, 47^. Therefore, future climate scenarios would enhance the competitive advantages and explain the increase of soil gram-positive pathogens, which include phytopathogens such as *Clavibacter michiganensis* and *Rhodococcus fascians* and human pathogens like *Staphylococcus aureus, Mycobacterium tuberculosis, Streptococcus pneumoniae* and *Listeria monocytogenes*. Among these soil-borne pathogens, *Clavibacter michiganensis* is an antibiotic-resistant phytopathogen that is subdivided into five subspecies according to their different host ranges, that is tomato, potato, maize, wheat and alfalfa ^48^. The *Staphylococcus aureus*, frequently detected as multi-antibiotic-resistant isolates, is a dangerous and versatile human pathogen that can cause skin and respiratory tract infections ^49^. In addition, we also found warmer temperatures increased the proportion of phytopathogenic fungi (e.g., *Verticillium dahliae*), which complies with a recent report on the increase in soil-borne plant fungal pathogens with elevated soil temperature ^50^.

In conclusion, our results highlight the significant impacts of future climate scenarios on soil resistome and pathogenome of cropland and grassland. Climate warming and extreme summers can significantly change the composition of microbial communities, leading to the enrichment of specific ARG types. Importantly, climate warming can promote the dissemination of ARGs, therefore elevating the overall health risk of soil resistome. Climate warming and extreme summers also disrupt the ecological balance of the soil microbiome, providing a competitive advantage for specific soil-dwelling plant or human pathogens. This escalation in the likelihood of outbreaks of particular plant and human diseases is poised to result in significant economic losses and pose a severe threat to human health worldwide. Overall, our findings emphasize the necessity of continuous monitoring of soil resistome and pathogens under the on-going global climate change, which is critical to protect human health and the sustainability of modern agriculture in a global One-Health framework.

## Data availability

The metagenome assemblies and metagenome-assembled genomes were deposited in the China National GeneBank (CNGB) database with the accession number CNP0005252.

## Authors’ contributions

F. J. obtained funding and supervised the research work. Z. Z. conceptualized this study, analyzed the data and wrote the manuscript. F. J. revised and finalized the manuscript.

## Acknowledgment

The authors thank Ms. Jiajing Guo for laboratory management support and Mr. Guoqing Zhang for the meaningful discussions and management of the laboratory computational server. The authors thank the Global Change Experimental Facility (GCEF) at Helmholtz-Centre for Environmental Research, Germany for making their metagenomic data (PRJNA 838942) publicly available to the scientific community. The authors thank the Westlake University HPC Center for computational support.

